# Cell adhesion strength and tractions are mechano-diagnostic features of cellular invasiveness

**DOI:** 10.1101/2021.12.30.474608

**Authors:** Neha Paddillaya, Kalyani Ingale, Chaitanya Gaikwad, Deepak Kumar Saini, Pramod Pullarkat, Paturu Kondaiah, Gautam I. Menon, Namrata Gundiah

## Abstract

The adhesion of cells to substrates occurs *via* integrin clustering and binding to the actin cytoskeleton. Oncogenes modify anchorage-dependent mechanisms in cells during cancer progression. Fluid shear devices provide a label-free, non-invasive way to characterize cell-substrate interactions and heterogeneities in the cell populations. We quantified the critical adhesion strengths of MCF7, MDAMB-231, A549, HPL1D, HeLa, and NIH3T3 cells using a custom fluid shear device. The detachment response was sigmoidal for each cell type. A549 and MDAMB-231 cells had significantly lower adhesion strengths at τ_50_ than their non-invasive counterparts, HPL1D and MCF7. Detachment dynamics was inversely correlated with cell invasion potentials. A theoretical model, based on τ_50_ values and the distribution of cell areas on substrates, provided good fits to data from de-adhesion experiments. Quantification of cell tractions, using the Reg-FTTC method on 10 kPa polyacrylamide gels, showed highest values for invasive, MDAMB-231 and A549, cells compared to non-invasive cells. Immunofluorescence studies show differences in vinculin distributions: non-invasive cells have distinct vinculin puncta, whereas invasive cells have more dispersed distributions. The cytoskeleton in non-invasive cells was devoid of well-developed stress fibers, and had thicker cortical actin bundles in the boundary. These correlations in adhesion strengths with cell invasiveness, demonstrated here, may be useful in cancer diagnostics and other pathologies featuring misregulation in adhesion.

## Introduction

Cell adhesions to substrates occur *via* integrin clustering. The binding of integrins to the actin cytoskeleton regulates essential cellular processes such as spreading, generation of contractility, migration, and cell cycle progression [1,2]. Oncogenes modify adhesion mechanisms during cancer progression, resulting in complex molecular cascades that alter cell migration, proliferation, and invasion [3–5]. Changes to the cell-cell adhesivity and E-cadherin abrogation are associated with the metastatic phenotype [6,7]. Cell stiffness correlates with metastatic and invasiveness potentials, with transformed and metastatic cells demonstrating increased deformability as compared to non-invasive cells [8,9]. Focal adhesions, including integrins and other structural and signaling proteins, are altered and more dynamic in invasive cancer cells [7,10,11]. Integrin antagonists have been used in clinical trials to block cell proliferation, survival, and migration, as well as progression and metastasis [12]. Being able to measure the strength of cell adhesions to substrates and tractions is potentially helpful to evaluate cell adhesion, with applications in targeted drug development for cancer. Such techniques also show promise in mechano -diagnostics, where sorting cells with differential adhesion strengths from a heterogeneous population of tumors might be required.

Spinning disc devices use hydrodynamic shear stress, in the range of 0-100 Pa on cell-seeded substrates, to quantify the strength of adherent cells [5,13,14]. Because shear stresses vary along the radial direction in such studies, cells are subjected to differing stresses based on their location on the substrate. Experiments using spinning devices show exponential variation in the adhesive strengths with bond clusters [13,14]. More recent studies demonstrate heterogeneities in the adhesivity of cancer cells; strongly adherent cells were less migratory than metastatic cells [15]. Microscope mounted, cone plate devices with small cone angles, exert uniform fluid shear stress on cell monolayers [16,17]. Such devices also present advantages in permitting real-time visualization of the stress fibers and focal adhesions under physiologically relevant and controlled shear stress conditions (0-10 Pa). Earlier studies using such a device showed that the number of cells on the substrate decreased in a sigmoidal fashion with an increase in shear stress [17]. A theoretical model, incorporating stochasticity of cell adhesions to the substrate and population-level differences in cell sizes, was able to recapitulate the experimentally obtained detachment curves [17]. These studies suggested the importance of the adhesive areas in the critical shear stress required to detach cells from substrates. Other studies delineated weakly adhered cells from those with strong adhesions, in the presence of Mg^+2^ and Ca^+2^ ions, using parallel plate flow chambers [18]. Highly metastatic cells had weak adhesions and were characterized by the disassembling of focal adhesions [19]. Traction studies demonstrate that metastatic cells are significantly more contractile and have dynamic focal adhesions as compared to non-metastatic cells that have stronger adhesions [19,20]. Focal adhesions and stress fiber contractility are both important parameters that contribute to the cell adhesions to substrates [20].

We use a custom fluid shear device to obtain de-adhesion curves for cancer cells with differential invasiveness, including breast epithelial cells (MCF-7 and MDAMB-231), lung epithelial cells (A549 and HPL1D), HeLa, and NIH3T3 fibroblasts. We fit experimentally obtained de-adhesion data, and cell area distributions to the theoretical model developed by Maan and co-workers [17]. Results show that invasive cells have significantly lower detachment strengths as compared to the less invasive cells in our study. Cell adhesive areas increased with invasion potentials. The theoretical model was able to capture the sigmoidal detachment curves for all cell types in the study. Invasive cells also had higher tractions, obtained using a regularized FTTC approach, as compared to other cells [21]. A quantification of the cell-substrate adhesion strength is important in diseases, such as cancer, featuring misregulation in adhesion.

## Methods

### Cell culture

MCF7, MDAMB-231, HPL1D [22], A549, Hela, and NIH3T3 cells in early passage were maintained in Dulbecco’s Modified Eagle Medium (DMEM, Invitrogen) supplemented with 10% (v/v) fetal bovine serum (FBS, Invitrogen) and 1% (v/v) penicillin/streptomycin (Sigma-Aldrich) in a humidified incubator containing 5% CO_2_ at 37°C. Cells were passaged every 2–3 days during the study.

### Measurement of the cell areas and immunofluorescence analysis

Cells were cultured on 22 mm coverslips, rinsed twice with chilled phosphate-buffered saline (PBS, Sigma Aldrich, D5652), and fixed with 4% paraformaldehyde solution for 15 minutes. To quantify the adhesive areas, cells were stained using rhodamine-phalloidin (1:200, Invitrogen, R415) at room temperature in antibody staining buffer. These data were used to obtain the cell areas (N∼100) for each group using Fiji (ImageJ). Data from adhesive areas in each cell group were fit to a log-normal distribution to obtain the mean, µ, and standard deviation of logarithmic values, s, for the distributions.

Specimens were permeabilized with 4% paraformaldehyde and incubated for 1 hour at room temperature in primary antibody anti-vinculin (1:500, Sigma Aldrich, V4505), secondary antibody Alexa Fluor 488 (1:600, Invitrogen, A32723), and rhodamine-phalloidin (1:200, Invitrogen, R415) at room temperature in antibody staining buffer to visualize stress fibers and vinculin. Cell nuclei were labelled with DAPI (1:500, Thermo, 62248) for 2 minutes, and the samples were rinsed thrice with PBS. Specimens were fixed with ProLong™ Diamond Antifade Mountant (Invitrogen, P36961), and imaged using a confocal microscope (Leica SP8, Bioimaging facility, IISc).

### Fluid shear experiments using the microscope mounted device

Glass coverslips (22 mm, Bluestar) were cleaned, air-dried, and attached to a 60 mm milled petri plate base using a thin layer of vacuum grease. Coverslips were plasma-activated for 2 minutes and coated with 40 µg/ml collagen I (Gibco) at 37°C for 1 hour in a humid chamber. Substrates were washed three times with phosphate-buffered saline (PBS), and cells were seeded at 5000 cells/ml for 12 hours to permit attachment and spreading. Hoechst (1:400; Thermo Scientific) was used to stain the nuclei for 3 minutes before the de-adhesion experiments. The cell-seeded petri dish was placed on the base plate of the fluid shear device on an inverted microscope (Leica DMI6000B with PeCon incubator) maintained at 5% CO_2_ at 37°C (Fig. 1a). The device works broadly on the principle of a cone-plate rheometer, and subject’s cells to Couette flow using media (Fig. 1b) [17]. A conical disc with 1° cone angle was attached to a motor, extracted from a computer hard drive, and driven using an Electronic Speed Controller (ESC). The device was powered using a DC power supply, and an Arduino UNO circuit was used to control the pulse width modulation signal [23]. The cone was rotated at different speeds using a custom program, and the speed of rotation, *ω*, was determined using a tachometer. Levelling screws on the device were used to align the cone parallel to the petri dish. The cone was positioned ∼10 µm above the petri dish base using a translation stage, and the fluid shear stress was increased in 0.2 Pa steps each minute using the ESC (Fig. 1c). Shear stress, *τ*, was calculated using the fluid viscosity, *μ*, cone angle, *β*, and rotation, *ω*, as

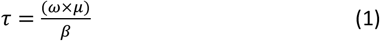

**Figure 1.**
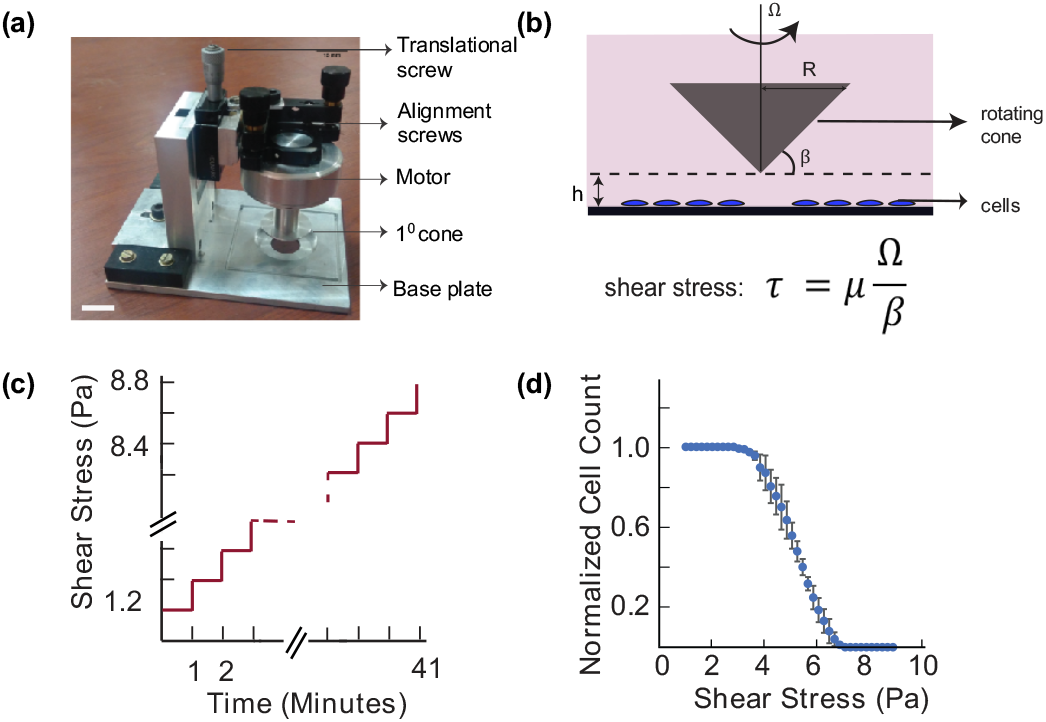
**(a):** Image shows a custom-fabricated, microscope mountable, fluid shear device used in the study. Device parts are labelled and include a 1° cone which is rotated using a motor on a petri dish containing adherent cells. The motor is aligned such that the cone is orthogonal to the plate. A translational stage helps position the cone at a desired height relative to the base plate. Scale Bar=10mm. **(b)** Couette flows are generated through rotation of the cone using a controlled program to exert shear stresses on the cell monolayer. **(c)** Shear stresses were increased in steps of 0.2 Pa at each minute, and images of the cells were taken using the inverted microscope. **(d)** The normalized number of cells remaining on the plate are plotted with each step increase of shear stress for a representative HeLa sample.

Images of cells, acquired during each increase of shear stress, were analyzed using Fiji (ImageJ) to obtain the number of cells remaining on the substrate. The normalized number of cells during each increase in shear stress was plotted to obtain the detachment response of cells (Fig. 1d).

Experiments were performed in biological triplicates, consisting of ∼100 cells for every run, for each of the six cell types in the study. Data are reported as mean ± SD. The cell detachment stress was compared between the various cell groups using a one-way analysis of variance (ANOVA) with Bonferroni comparisons to test for individual differences between the groups. Statistical differences (p < 0.05) are indicated in the plots.

### Theoretical model for deadhesions of attached cells under fluid shear

Experimentally obtained data were used in the theoretical model developed for cell detachments under fluid shear by Maan and co-workers [17]. In this model, the cell was approximated to be a solid hemisphere, and was assumed to have a uniform distribution of focal adhesions along the perimeter. The model cell was subjected to Stokes flow, characterized by low Reynolds numbers, and the kinetics of adhesions to the substrate were described using the Bell’s model. Based on this model, the attachments between a cell and the substrate form at a rate, k_on_, and break exponentially with the applied stress [24]. The number of cells remaining on the substrate, Φ(t), under a controlled increase in the fluid shear stress, τ, is given by

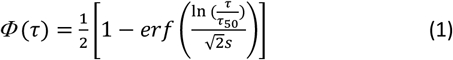

*τ*_50_ is the critical shear stress corresponding to 50% cells remaining on the substrate during increase in the fluid shear, and is obtained from the detachment curve. The parameter, s, in equation (1) was estimated using the best fit to cell area distributions as described earlier. This value is hence proportional to the areal width of the distributions.

### Traction force microscopy

Cleaned coverslips (22 mm) were treated at room temperature with 3-amino-propyl triethoxy silane (Sigma Aldrich) and incubated with 0.5% glutaraldehyde solution (SDFC Ltd.) for 30 minutes. Another set of cleaned coverslips was treated with poly-L-lysine (Sigma Aldrich) for 45 minutes at 37°C in a humid chamber. A 18 µl/ml of 2% stock fluorescent beads (Invitrogen, F8810) was spin-coated on the coverslip using a previously published protocol [21]. Solutions of acrylamide (PA; 40% wt/vol, Sigma Aldrich) and N, N’-methylene bis-acrylamide (2% wt/vol, Sigma Aldrich) were mixed with distilled water [25]. 30 μl of the solution was combined with 10% APS (Thermo, 17874) and TEMED (Thermo, 17919), sandwiched between the cleaned and bead coated coverslips, and polymerized at room temperature for 30 minutes to obtain gels of 10 kPa stiffness. The bead-coated coverslip was carefully removed to obtain the polyacrylamide gel, which was next attached to a 35 mm punched petri dish (Nunc, Thermo Scientific) using a thin layer of vacuum grease (SDFC Ltd.). A solution of heterobifunctional sulpho-SANPAH linker (200 µl of 100 mg/ml stock; Thermo, A35395), diluted in HEPES buffer (50 mM; Sigma Aldrich, H3375), was pipetted to the gel surface. The assembly was exposed to 365 nm UV light (Thermo, 95034) for 10 minutes. Collagen-I (100µl of 40 μg/ml concentration) was incubated on the gel surface at 37°C for 45 minutes in a humid chamber.

Substrates were rinsed thrice with HEPES buffer, and 200 ml of cells (2000 cells/ml) was seeded on the gels for 12 hours. The experiment was performed in a live cell chamber at 37°C and 5% CO_2_ (Leica DMI6000B with PeCon incubator; 40X oil immersion objective). The attached cells were used to quantify the traction stresses. Three images were obtained for each cell; first, a phase-contrast image to obtain the contour. Second, a fluorescent image of beads on the gel surface with the attached cell (stressed configuration), and finally, an image of the beads (referential configuration) following trypsinization (Sigma Aldrich). These images were used to quantify the constrained traction stresses exerted by cells on the substrate using a regularized Fourier Transform Traction Cytometry (Reg-FTTC) method in MATLAB [21]. Experiments were performed in triplicates for each of the different cell types in the study. Results from n=10 are reported for each cell group in the study.

## Results and Discussion

### Higher cell invasiveness decreases the cell adhesion strengths

We used sufficiently spaced cells on the petri dish to minimize the contributions from cell-cell interactions in the analyses. The fluid shear stress was ramped, and the number of cells remaining on the substrate was measured using the device for each of the six different cell types in the study (Fig. 1a). These include an embryonic mouse fibroblast (NIH3T3), human cancer cells of varying invasion potentials, and non-invasive cells. The HPL1D is an immortalized non-invasive lung epithelial cell line, whereas A549 cells are invasive lung adenocarcinoma cells. MCF7 is a non-invasive breast cancer cell line compared to the invasive MDAMB-231 cells. In addition, we also used HeLa cells which are metastatic cervical cancer cells in the study.

The cell detachment responses were sigmoidal for all cell types; the shapes and shifts of the detachment curves were however significantly different for all groups in the study. We delineated the detachment responses based on three different magnitudes of shear values (Fig. 2a). τ_10_ is the threshold shear stress, required to detach 10% of cells from the substrate, which helps identify loosely or weakly adhered cells on the substrate. τ_50_ is the characteristic critical de-adhesion strength based on removal of 50% of cells from the substrate under shear. Finally, τ_90_ shows the percentage of cells remaining on the substrate under fluid shear. This value indicates the ability of cells to reinforce focal adhesions and remodel in response to shear stress. We compared the detachment strengths, obtained from experiments at τ_10_, τ_50_ and τ_90_, with the invasive potentials of cells in the study. More invasive cells, A549 and MDAMB-231, had significantly lower critical detachment strengths, τ_50_, as compared to the non-invasive MCF7 (Figure 2b; Table 1). HeLa cells had significantly higher τ_50_ values than the MDAMB-231 and A549 cells (Fig. 2b, Table 1).

**Table 1:** Critical deadhesion strengths (τ_50_) obtained from the experiments and cell area distributions parameters (n=100 each group) from the log-normal distribution (µ,s) were used in the theoretical model to quantify cell detachment from the substrate under fluid shear.

| | $\tau_{50}$ (Pa)<br>(Mean ± SD) | Cell Area<br>($\mu\text{m}^2$)<br>(Mean ± SD) | $\mu$ | s | $\chi^2$ |
| --- | --- | --- | --- | --- | --- |
| NIH3T3 | $5.1 \pm 0.10$ | $1928.06 \pm 481.79$ | 7.54 | 0.25 | 3.73 |
| HPL1D | $4.27 \pm 0.15$ | $1866.76 \pm 452.11$ | 7.50 | 0.24 | 2.24 |
| MCF-7 | $4.70 \pm 0.10$ | $757.71 \pm 151.22$ | 6.61 | 0.21 | 1.65 |
| HeLa | $4.63 \pm 0.35$ | $880.95 \pm 162.44$ | 6.76 | 0.19 | 1.32 |
| A549 | $3.20 \pm 0.10$ | $1049.68 \pm 170.65$ | 6.94 | 0.16 | 2.93 |
| MDAM B-231 | $3.47 \pm 0.31$ | $1248.01 \pm 254.25$ | 7.11 | 0.20 | 6.16 |

**Figure 2.**
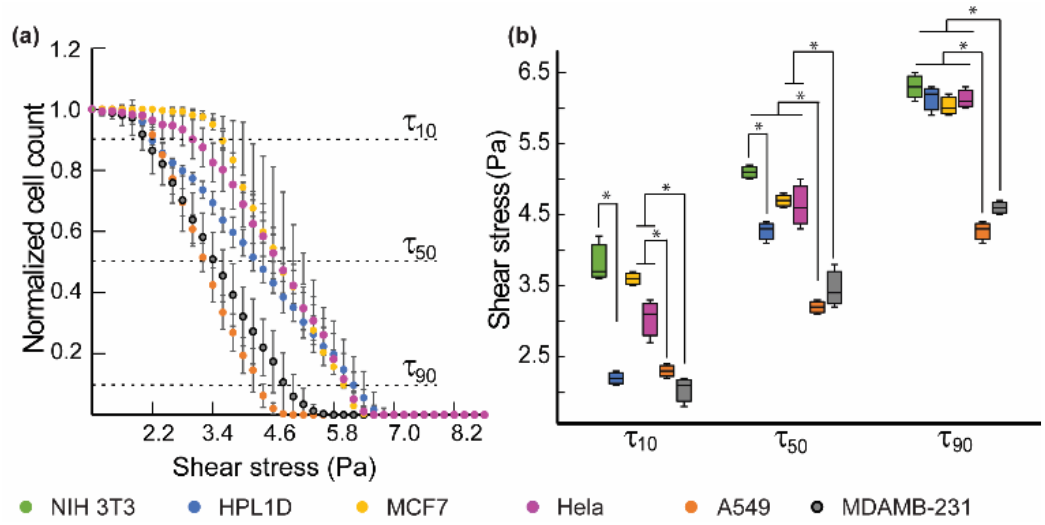
**(a):** Normalized cell count, calculated as the number of cells at a given time with respect to the number of cells at the start of the experiment, are plotted as a function of shear stress for each of the six different cell types in the study. Data are represented as Mean ± SD. **(b)** The de-adhesion strengths corresponding to τ_10_, τ_50_, and τ_90_ were compared between the different groups. Statistically significant differences (p<0.05) are indicated by (*).

Normal lung epithelial cells (HPL1D) had significantly higher τ_50_ values as compared to the more invasive (A549) cells. The NIH3T3 cells had the highest critical detachment strength in the study (Table 1). Critical detachment strength hence shows an inverse correlation with the invasive potential of cells. The threshold de-adhesion values of shear (τ_10_) were also clearly different in the various cell types in our study. Fibroblasts (NIH3T3) had the highest values of τ_10_ (3.83 ±0.32 Pa) whereas the metastatic cells (A549 and MDAMB-231) had significantly weaker adhesions characterized by low values of τ_10_. The non-invasive MCF7 cells had significantly higher values of τ_10_ as compared to the more invasive cells in the study. The HPL1D cells had lower τ_10_, and very high values of τ_90_, which suggests heterogeneity in the cell adhesion strengths. Cancer cells with higher invasive potentials, A549 and MDAMB-231, had significantly lower values of τ_90_ under sustained shear as compared to other cells. There were no differences in these values for all other cell groups.

A log-normal distribution provided good fits to the experimentally measured spread areas for all cell types in the study (Fig. 3; Table 1). MCF7, HeLa, and A549 cells had a narrow distribution of areas as compared to other cells in the study. NIH3T3 and HPL1D cells had higher mean spread areas and larger standard deviations. We plotted the mean cell areas and critical detachment strengths (τ_50_) to investigate possible correlations between these parameters (Fig. 4). Cell areas showed a linear increase with higher cell invasiveness potentials. Although the MCF7 and A549 cells had similar projected areas, the former had a significantly higher value of critical detachment strength (τ_50_) as compared to A549 cells. Comparisons between the MCF7 and MDAMB-231, and the HPL1D and A549 groups show that more invasive cells within the groups required lower critical shear stress to detach as compared to those with lower invasive potentials. The normal HPL1D cells required higher critical shear stress to detach and had greater cell spread areas as compared to all other cancer epithelial cells in the group. NIH3T3 fibroblasts had the highest critical stress to detach from substrates (5.1±0.10 Pa), and the highest mean area in the study. These data suggest that a small population of strongly adherent cells, remaining under sustained shear, may be due to cell polarizations.

**Figure 3:**
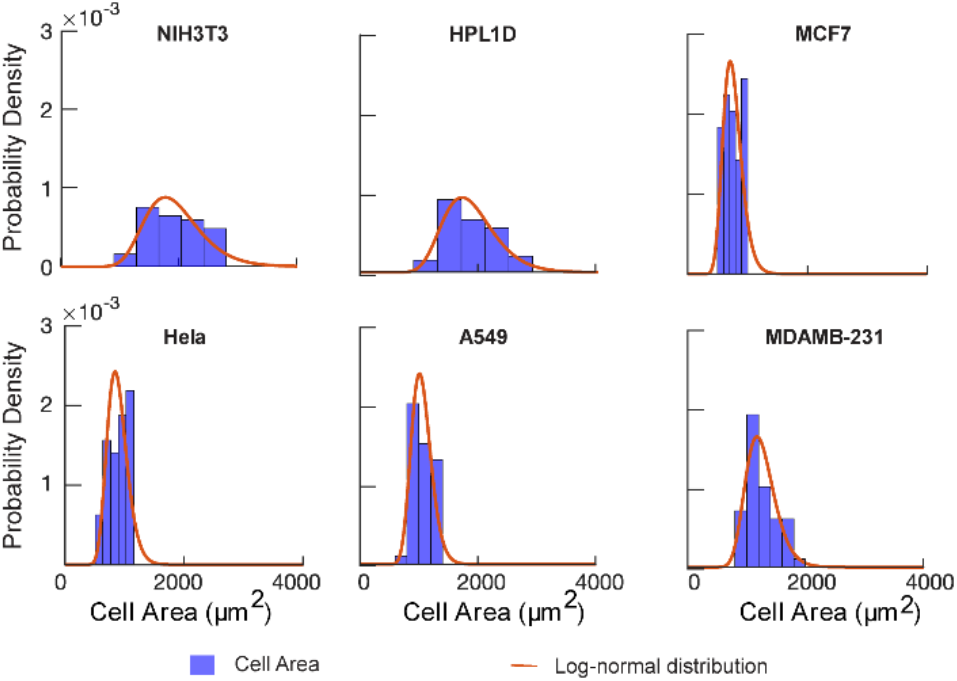
Cell spread areas for each group (n=100) were plotted based on the probability density, and the distributions were fit to a log-normal function.

**Figure 4:**
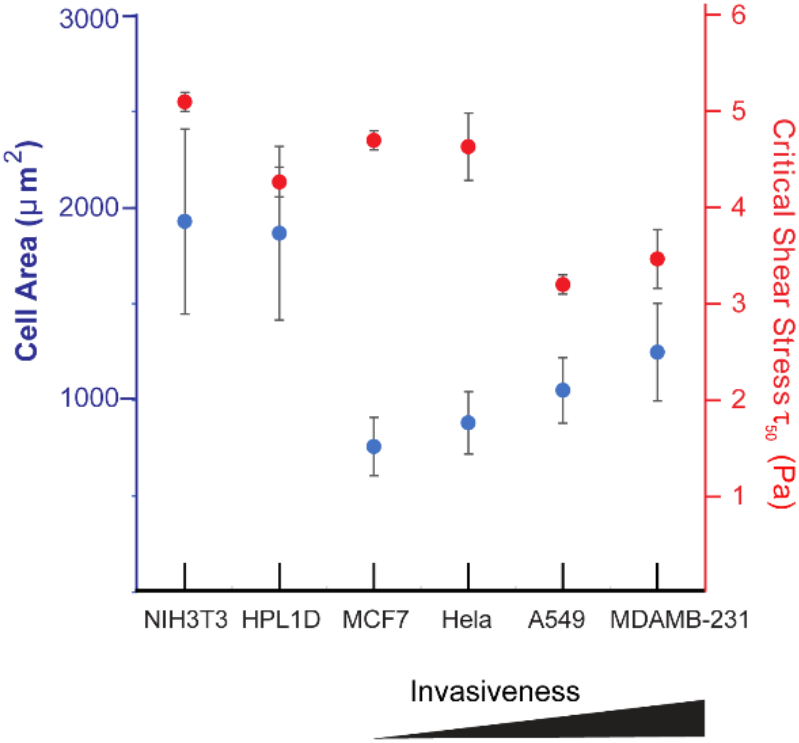
Cell areas and critical shear stress (τ_50_) are plotted based on the cell invasiveness.

The sigmoidal detachment profiles obtained in this study are similar to the results reported by Maan *et al*. for HEK and NIH3T3 cells using a similar device [17]. Other studies have also reported deadhesion profiles with marginally different critical detachment strengths for various cell types, including osteosarcoma cells, NIH3T3, WI38 fibroblast, Swiss 3T3 murine fibroblast, among others [13,18,26–28]. Cancer cells regulate cell-substrate interactions during the various stages of metastasis through changes in their adhesion kinetics [29]. The types of integrins in breast cancer cells (αvβ5/αvβ3 and α5β1/α2β1) also change under static and shear flow conditions [30]. Variations in the adhesion strengths of cancer cells may also depend on the cell type and the presence of oncogenes, such as ERBB2, induced during metastasis, and correlated with differences in the integrin types [31,32,33]. Less metastatic HT-29P cells had six times higher adhesion strengths than the highly metastatic HT-29LMM on collagen-1 substrates; these adhesions were however not different on fibronectin-coated substrates [34]. Marginal differences in the critical detachment strengths of cells in our study to those reported earlier may be due to the differences in the extracellular matrix type, ligand densities, and the application of shear stress using media in this study as compared to the use of PBS alone by others [16,18,28]. Cell detachment studies hence provide a quantitative measure of the differences in adhesion kinetics between cells of varied invasive potential.

### Modelling cell detachments from substrates using τ_50_ and cell adhesive areas

We used the experimentally measured critical de-adhesion strengths (τ_50_) and adhesive areas of cells on ligand coated substrates (Fig. 3) to test model predictions for the various cell types in our study (Fig. 5). The model showed overall good fits to the experimentally obtained results for all cells in our study. De-adhesion data for the MCF7 cells fit the model predictions very well. The model, however, marginally overpredicts the results for the HPL1D cells at τ_90_. Variations in the model fits at τ_90_ are also apparent for MDAMB-231 and NIH3T3 cells which have elongated morphologies that deviate significantly from the spherical shape assumed in the model. Cell projected areas have inherent heterogeneities at the tails of the distribution that may also contribute to the differences with the model predictions. Deviations from the model may also be related to possible cell remodeling and reinforcements in the adhesions under sustained shear [35]. In contrast, τ_10_ values were marginally overpredicted by the model, as compared to experimentally obtained values, for all cell types barring the MCF7 and HPL1D cells. These differences may either be due to weak adhesions or possible biological heterogeneities within the cell populations which we have not assessed in this study.

**Figure 5:**
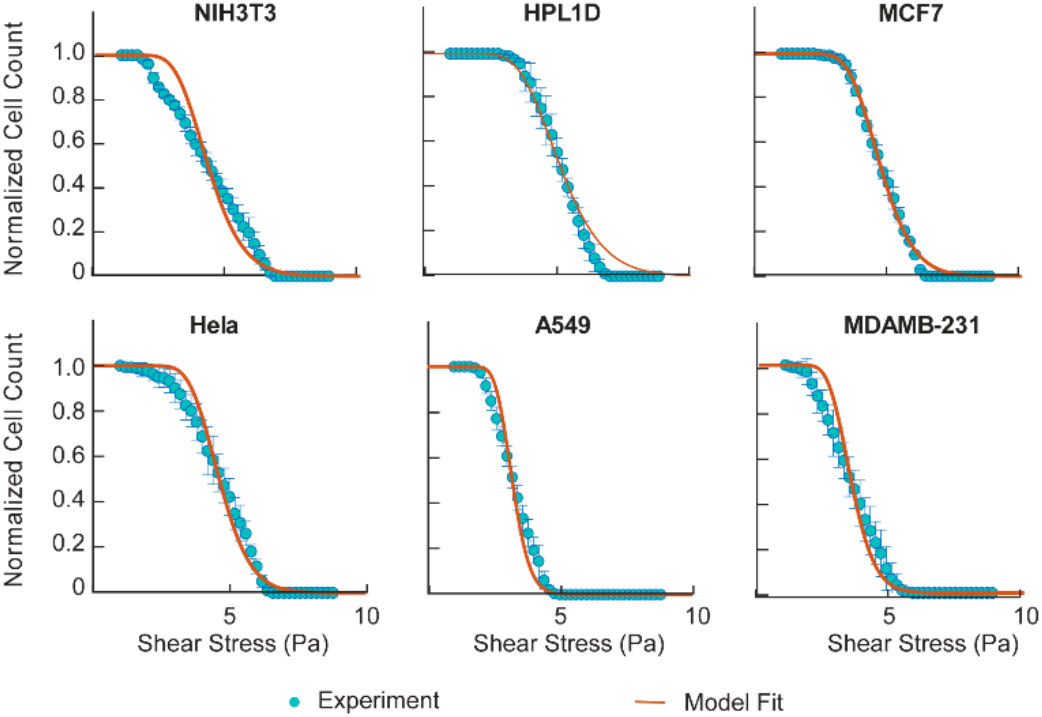
Model predictions show good fits with experimental data for all cell types. The model uses τ_50_ values, and the mean, µ, and SD, s, values from the areal distributions for each cell type in the study.

Cells form integrin-mediated adhesions with the substrate at the leading edge, and disintegrate at the retracting end during migrations. Adhesions strengthen with cell spreading and result in an increase in the cell-substrate contact area, receptor clustering, and focal adhesion assembly, through interactions with the cytoskeleton [36] and the bound integrins [5]. A higher ECM ligand density also controls cell spreading through focal adhesion assembly. Cell spreading regulates function through changes in the morphology and cytoskeletal tension [37,38]. Modifications to the model, including variations in the cell shape under shear, possible redistributions in focal adhesions due to cell polarizations, and varying stress fiber contractility, may be useful in future studies to better estimate the marginal deviations in the detachment profiles of cells from substrates. The theoretical model, based on τ_50_ and the adhesive areas of contact between the cell and the substrate, is however useful to delineate the differences in critical de-adhesion strengths between cell types.

### Invasive cells exert higher traction stresses

We used traction force microscopy to quantify the differences in cell contractility in the different cell types (Table 2) using 10 kPa polyacrylamide gels (n=10 in each group). Fig. 6a shows the maximum tractions exerted by adherent cells on substrates obtained using the Reg-FTTC approach [21]. There were no statistical differences in maximum tractions between MCF7 (294.45±48.74 Pa) and HeLa cells (293.58±36.32 Pa) that had the lowest tractions among all cells in the study (Table 2). In contrast, the MDAMB-231 cells had the highest tractions (2050.82±127.29 Pa), followed by the A549 invasive cancer cells (1116.24±86.71 Pa). NIH3T3 fibroblasts and normal lung epithelial cells (HPL1D) had intermediate tractions (833.12±92.01 Pa and 820.58±186.81 Pa, respectively), which were higher than the non-metastatic cells, and lower than corresponding metastatic cells. The Reg-FTTC approach uses a regularization parameter (γ*) which is determined using an inflection point in the plots of the maximum tractions with the log of the regularization parameter [21]. Although the maximum tractions obtained using Reg-FTTC were lower than those obtained using the FTTC approach, similar trends were visible for the different cell types in this study. The spread areas for the different cell types were next plotted to explore possible correlations with tractions. Cells with greater invasiveness potentials exerted higher traction stresses on 10 kPa gels as compared to cells with lower invasion potentials in our study.

**Table 2:** Results from traction force microscopy (FTTC) are shown for all cells in the study (Mean±SD). Values obtained from constrained and regularized approach (Reg-FTTC) are presented, and the range of regularization parameters are indicated (n-10 each cell type).

| | Max traction (FTTC) (Pa) | Max traction (Reg-FTTC) (Pa) | Regularization parameter ($\gamma^*$) Range |
| --- | --- | --- | --- |
| NIH3T3 | 1032.14<br>$\pm 290.95$ | 833.12<br>$\pm 215.01$ | 2.053 to<br>2.244E-11 |
| HPL1D | 1017.29<br>$\pm 590.73$ | 820.58<br>$\pm 483.65$ | 2.054 to<br>2.539E-11 |
| MCF7 | 358.22<br>$\pm 114.87$ | 293.58<br>$\pm 102.84$ | 4.071 to<br>5.411E-11 |
| HeLa | 372.76<br>$\pm 154.13$ | 294.45<br>$\pm 125.38$ | 1.572 to<br>1.573E-10 |
| A549 | 1398.07<br>$\pm 274.19$ | 1116.24<br>$\pm 228.94$ | 4.067 to<br>1.13E-10 |
| MDAMB-231 | 2628.62<br>$\pm 402.53$ | 2050.82<br>$\pm 308.33$ | 1.443 to<br>5.411E-11 |

**Figure 6:**
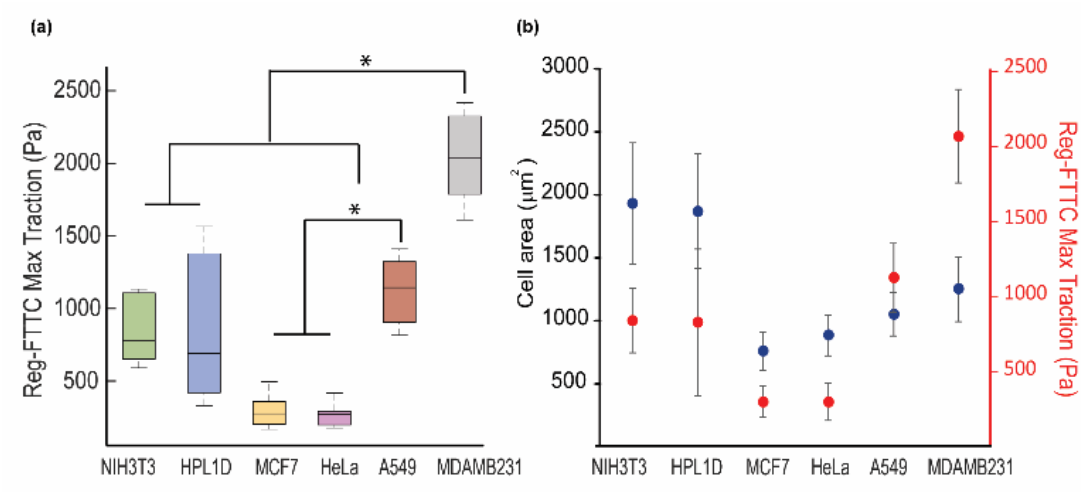
**(a)** Cell tractions were measured using the Reg-FTTC method and are plotted as Mean ± SD in the study (n=10 each group). **(b)** Cell areas and Reg-FTTC tractions increase with invasiveness potential. Statistically significant differences (p<0.05) are indicated by (*).

These data agree with earlier studies which show that metastatic cells exert higher traction stress as compared to non-metastatic cells [20]. Tumor-associated endothelial cells also exert higher tractions than normal endothelial cells [39]. In contrast, murine breast cancer cells with higher metastatic potential, and Ras-transformed fibroblasts, showed weak traction stresses [10,40]. H-ras transformed 3T3 fibroblasts exert lower tractions than normal 3T3 fibroblasts cells [41]. Lung and breast cancer cells with higher invasion potentials generate higher 3D tractions as compared to their counterparts with lower invasive potentials [42]. Focal adhesions and stress fibers are essential components in traction generation by adherent cells on substrates. The traction stresses exerted on substrates may hence be used as a biophysical marker of the metastatic potential and malignancy of cells [20,42,43].

### The cytoskeleton and invasiveness potential of cells

Faster cell detachment in cell types with higher invasion potential suggests a possible role for cell contractility, and focal adhesions. We stained for actin and vinculin distributions to visualize differences in the stress fibers and focal adhesions among the different cell types in the study. Immunofluorescence studies show apparent differences in the expression of vinculin in the various cell types (Fig. 7). Cells with lower invasion potentials had distinct and well-formed vinculin puncta that are reminiscent of stationary and less migratory cells. The cytoskeleton in these cells also have thick cortical actin bundles, and lack well-developed stress fibers (Fig. 8). In contrast, cells with higher invasion potential had diffuse vinculin. Lamellipodia and protrusions are clearly visible in the migratory cancer cells that have higher invasion potentials.

**Figure 7:**
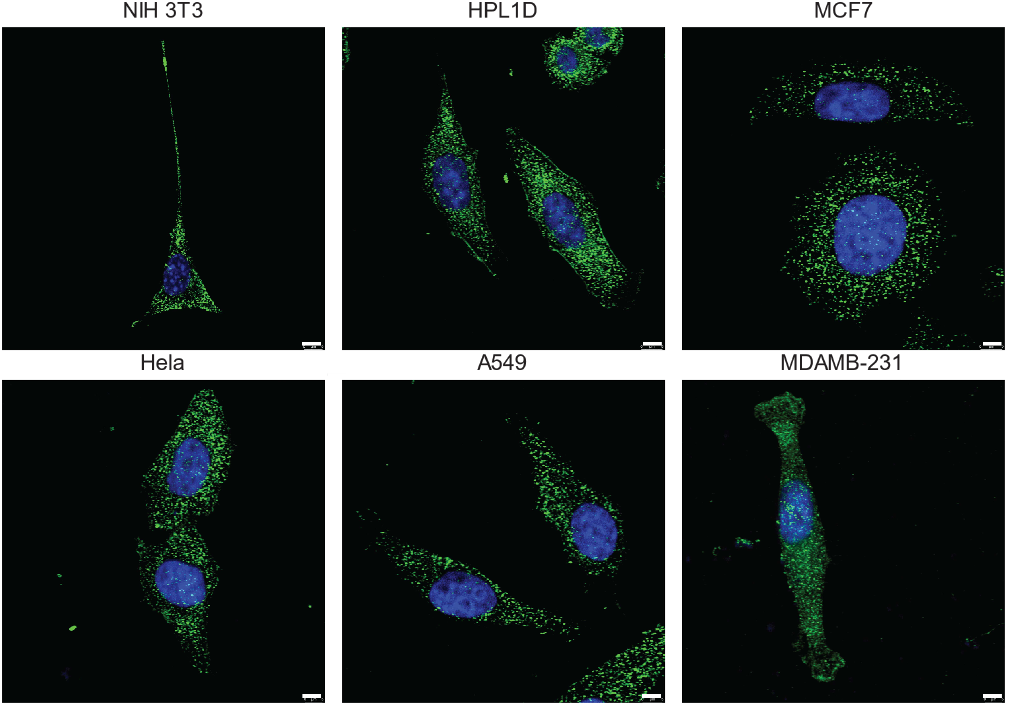
Focal adhesion sizes and distribution are visibly different and vary based on cell invasiveness. Vinculin in the focal adhesions is labelled using green, and the cell nuclei are in blue. Scale bar=10µm.

**Figure 8:**
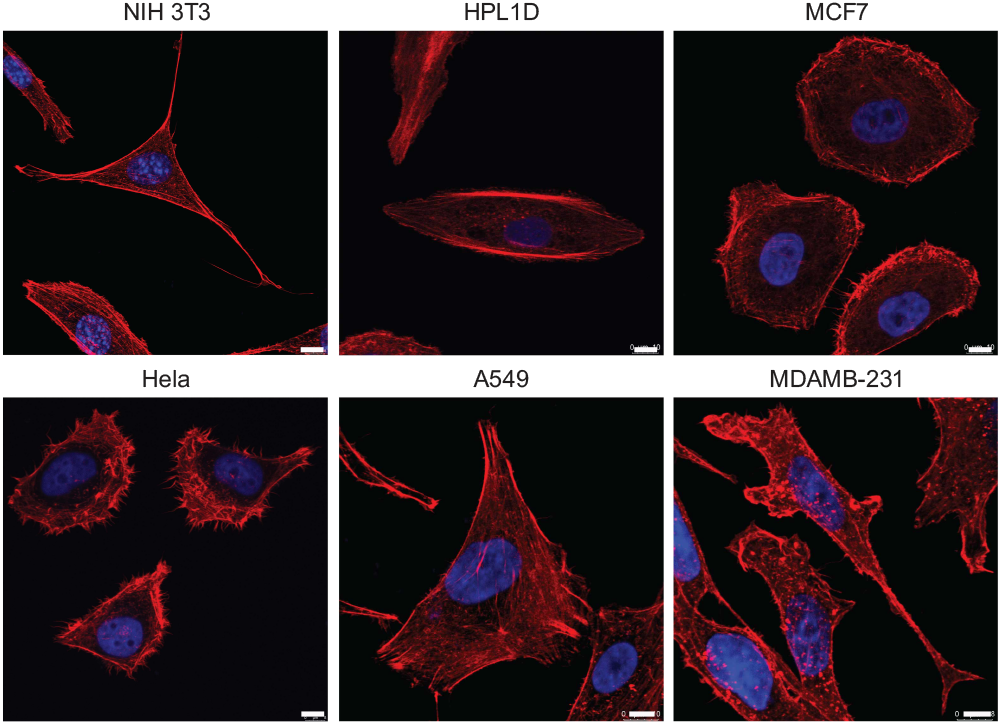
F-actin stained images of the cells show prominent stress fibers in A549 and MDAMB-231 cells. Non-invasive cells showed actin at the boundary and cortex. Actin is labelled red, and the nucleus is blue. Scale bar=10µm.

The differences in focal adhesion assembly, migration, and contractility determine the invasion potentials, which is linked to the downregulation of E-cadherin in cells that undergo the epithelial-to-mesenchymal transitions (EMT) [44]. Genetic and environmental cues cause changes to the adhesions during EMT, resulting in heterogeneities in the adhesive phenotype [45,46]. Beri *et al*. delineated weakly and strongly adherent cells using fluid shear from cell populations [47]. Heterogeneity in cancer cells may be genetic, epigenetic, transcriptomic, proteomic, and functional [48]. Cancer cells actively modify their spreading [49,50], and migration [51,52] behaviors in response to biophysical cues. Clonal heterogeneity analysis is useful to predict the patient prognosis and response to therapy. Fluid shear stress experiments may be useful in delineating heterogeneities in cell populations based on the cell adhesion strengths to substrates.

## Conclusions

Cancer metastasis is characterized by the local invasion of cells from primary tumors, their transport through the systemic circulation, their extravasation to secondary sites and the growth of metastatic lesions in those sites [53]. The invasive ability of cancer cells correlates with their ability to modulate their adhesion strengths. Experiments using a fluid shear device have proved useful to delineate differences in adhesion strengths between cancer cells, to sort cells based on differential adhesion abilities and to quantify the differences in the roles of adhesions during cancer metastasis.

Migration is inhibited in cells that are firmly adherent to the ECM [54]. We show that more metastatic cells have lower adhesion strength than non-metastatic cells, while normal cells have higher adhesion strength than all cancer cells. The theoretical model of Maan *et al*. provides reasonably good fits to the deadhesion profiles for all cell types in our study. A marginal deviation from the predicted sigmoidal behavior may arise from the differential contractility, assumptions regarding the FA size and distribution, and the differences in cell shapes that are not included in the model.

Non-invasive cells have a dense meshwork of peripheral actin filaments and relatively few stress fibers. Invasive cells have higher contractility as compared to non-invasive cells. Normal cells, however, exert higher tractions than invasive and non-invasive cells and have prominent stress fibers. Focal adhesions are more prominent in non-invasive cells as compared to invasive and normal cells. The adhesion strength and traction measurements are hence useful as biophysical markers of cell metastasis. Inherent differences in the differential adhesion strengths may help sort cells with varying invasive potentials. Additional experiments with inhibitors for proteins involved in cell-substrate mechanics and contractility will be useful in understanding the functional regulatory circuits that may be involved in these interactions and in the design of better therapeutic options.

## Author Contributions

NP performed all experiments, analyzed results, and helped write the manuscript. KI helped perform the traction experiments, and CG helped with the device design. DKS and PK provided inputs with cell culture and immunofluorescence studies. PP, GIM, and NG designed the study and analyzed the results. NG wrote the manuscript with inputs from all authors.

## Acknowledgements

We thank the imaging facility in Biological Sciences, IISc, for the confocal imaging reported in this study. We thank members of the mechanical workshop at Raman Research Institute for help with device fabrication. PK and DKS laboratories are supported by the IISc-DBT partnership program, and DST-FIST infrastructure to MRDG. PK is a recipient of senior scientist fellowship from the Indian National Science Academy. PP, GM, and NG are grateful to the Department of Biotechnology (BT/PR23724/BRB/10/1606/2017) for project support through a jointly funded grant.

